# Phylogeny-aware simulations suggest a low impact of unsampled lineages in the inference of gene flow during eukaryogenesis

**DOI:** 10.1101/2024.10.04.616067

**Authors:** Moisès Bernabeu, Saioa Manzano-Morales, Toni Gabaldón

## Abstract

Gene phylogenies are broadly used to analyse events of horizontal gene transfer, namely, their presence, the potential donor and acceptor lineages, and their relative timing. Recent phylogenomics analyses have reconstructed a relative chronology of gene acquisitions in the lineage leading to the eukaryotes, revealing waves of acquisition from different donors before and after the mitochondrial endosymbiosis. However, a recognised caveat is the potential biases introduced by the presence of incomplete taxon sampling resulting in so-called “ghost” lineages. Here, we assessed the robustness of the gene phylogeny-based branch length ratio method in the inference of the relative ordering of gene acquisition events during eukaryogenesis. We introduce a novel simulation framework that populates a known dated Tree of Life with plausible “ghost” lineages and simulates their gene transfers to the lineage leading to the last eukaryotic common ancestor. Our simulations suggest a generally low probability of misinterpreting the relative order of gene acquisitions from distinct ghost donors. However, we identify specific problematic phylogenetic placements where unsampled lineages are more likely to produce misleading results. Overall, our approach offers valuable guidance for the interpretation of future work on eukaryogenesis, and can be readily adapted to other evolutionary scenarios.

## Introduction

Processes of non-vertical evolution, such as horizontal gene transfer (HGT) or hybridization are major drivers of genome evolution in all life domains (Arnold et al. 2021; Gophna & Altman-Price 2022; Gabaldón 2020; Douglas & Langille 2019). Beyond the topology, the length of the branches in a gene phylogeny can also convey information, typically about sequence divergence expressed by the number of residue substitutions per site. This divergence is the result of the combined effect of the time elapsed between the two nodes that delimit that branch, and the evolutionary rate at which that molecule has evolved in this period. In 2016, Pittis and Gabaldón proposed a “branch-length ratio method” to estimate differences in elapsed times between different groups of gene trees (Pittis & Gabaldón 2016a). This approach rests on the assumption that the ratio between the interrogated and normalising branches removes the rate component of the interrogated branch, obtaining its relative age. This framework was employed, among other studies, to reconstruct the relative timing of gene acquisition events during eukaryogenesis (Pittis & Gabaldón 2016a; Vosseberg et al. 2021). The conditions under which this approach can be applied were assessed and were shown to include those used in the original 2016 paper (Susko et al. 2021).

The approach has also triggered some methodological debates (William F. et al. 2017; Tricou et al. 2022; Susko et al. 2021; Bernabeu et al. 2024; Pittis & Gabaldón 2016b). One recent criticism (Tricou et al. 2022) has highlighted the potential for unsampled or extinct lineages (so-called “ghost lineages”) to introduce bias into phylogenetic analyses, particularly regarding the inference of the relative order of HGT events (Tricou et al. 2022). However, the simulations used to evaluate this potential had some relevant limitations (Bernabeu et al. 2024; Tricou et al. 2024), most importantly the use of simulated topologies that were entirely disconnected from the current knowledge on the tree of life and the relevant lineages in eukaryogenesis.

In this work, we aim to specifically test the potential impact of ghost lineages in the debate surrounding the relative order of events during eukaryogenesis (Pittis & Gabaldón 2016a; Vosseberg et al. 2021). In contrast to previous simulations (Tricou et al. 2022), here, we introduce a novel simulation framework that is grounded in two alternative recently reconstructed dated Trees of Life (Mahendrarajah et al. 2023; Moody et al. 2024) and that explicitly models the transfer of any potential ghost lineage branching out from currently known bacterial lineages. In this way, our model is able not only to assess the impact of ghost lineages in a specific phylogenomic analysis but also to identify the impact of alternative topologies and point at particularly problematic lineages or transfer scenarios. Beyond the study of gene transfers predating the origin of eukaryotes, our approach will certainly be useful to assess the potential impact of ghost lineages in other evolutionary questions.

## Methods

### Phylogeny-aware simulation framework

Given a dated (or ultrametric) species phylogeny and an acceptor branch representing the lineage that received the gene transfers, we simulate two ghost lineages in each iteration. These two lineages originate from branches in the species phylogeny that are different from the acceptor branch but that overlap, at least partially, with it. Then, for each simulated ghost we simulate a gene transfer to the acceptor branch at a randomly chosen time at which both ghost and acceptor branches exist. Assuming an inference is subsequently done in the absence of the ghost, the expected inferred donor lineage would correspond to the closest existing lineage, and the inferred length of the branch corresponding to the transfer will correspond to the time elapsed between the last common ancestor of the acceptor branch and the inferred donor, and the end of the acceptor branch. This allows us to assess differences between the true and the inferred order of gene transfer events, depending on the phylogenetic position of the putative ghost donors. The procedure is repeated for the desired number of iterations, and the species tree, the acceptor branch, and the number of ghost lineages can be varied depending on the scenario to test.

### Testing gene flow in eukaryogenesis

We adapted the above framework to specifically test the effect of ghosts on the scenario relevant for eukaryogenesis (Pittis & Gabaldón 2016a; Vosseberg et al. 2021). For this, we used as a species tree a recently inferred dated phylogenetic tree of life obtained from a concatenation of the ribosomal markers for the ATPase distribution analysis (Mahendrarajah et al. 2023) (the ATPase tree hereafter, Fig. 1a). We removed the chloroplast and mitochondria clades to obtain a reliable set of putative bacterial sisters for the ghost donor. We set the acceptor branch to be the branch connecting the last common ancestor of eukaryotes and their closest Asgardarcheota relatives (wich, for simplicity, we here name the first eukaryotic common ancestor (FECA), and the last eukaryotic common ancestor (LECA), the FECA-LECA lineage (Gabaldón 2021). We dated FECA and LECA using the tree branches (i.e., the mean of the posterior distribution of the node datings), obtaining a FECA-LECA time frame of 2.42 Ga to 1.89 Ga (Fig. 1b). Finally, we retrieved the birth and death of the branches present in the inferred FECA-LECA period, which we called the FECA-LECA branch space (Supplementary Fig. S1b). The simulated ghost lineages had to comply with the following conditions: (i) they branch out from a bacterial lineage (i.e. they are bacterial); (ii) they must have branched out from a branch that coexisted with the FECA-LECA branch; (iii) the transfer from the donor ghost lineage had to have occurred during the FECA-LECA period; (iv) the death of the ghost branch had to occur after the transfer, at any point up to the present (Fig. 1c). Note that this last point is a formalism and does not take part in the assessment of shifts. At the end of the simulation, we calculated the proportion of iterations in which the early/late classification is reversed if the lineages are unsampled, that is, when there is a “shift” in the relative ordering of the observed transfers. This process is agnostic to whether the lineage is extinct (its death occurred in the past) or just unsampled (data on its existence is still missing).

**Fig 1.**
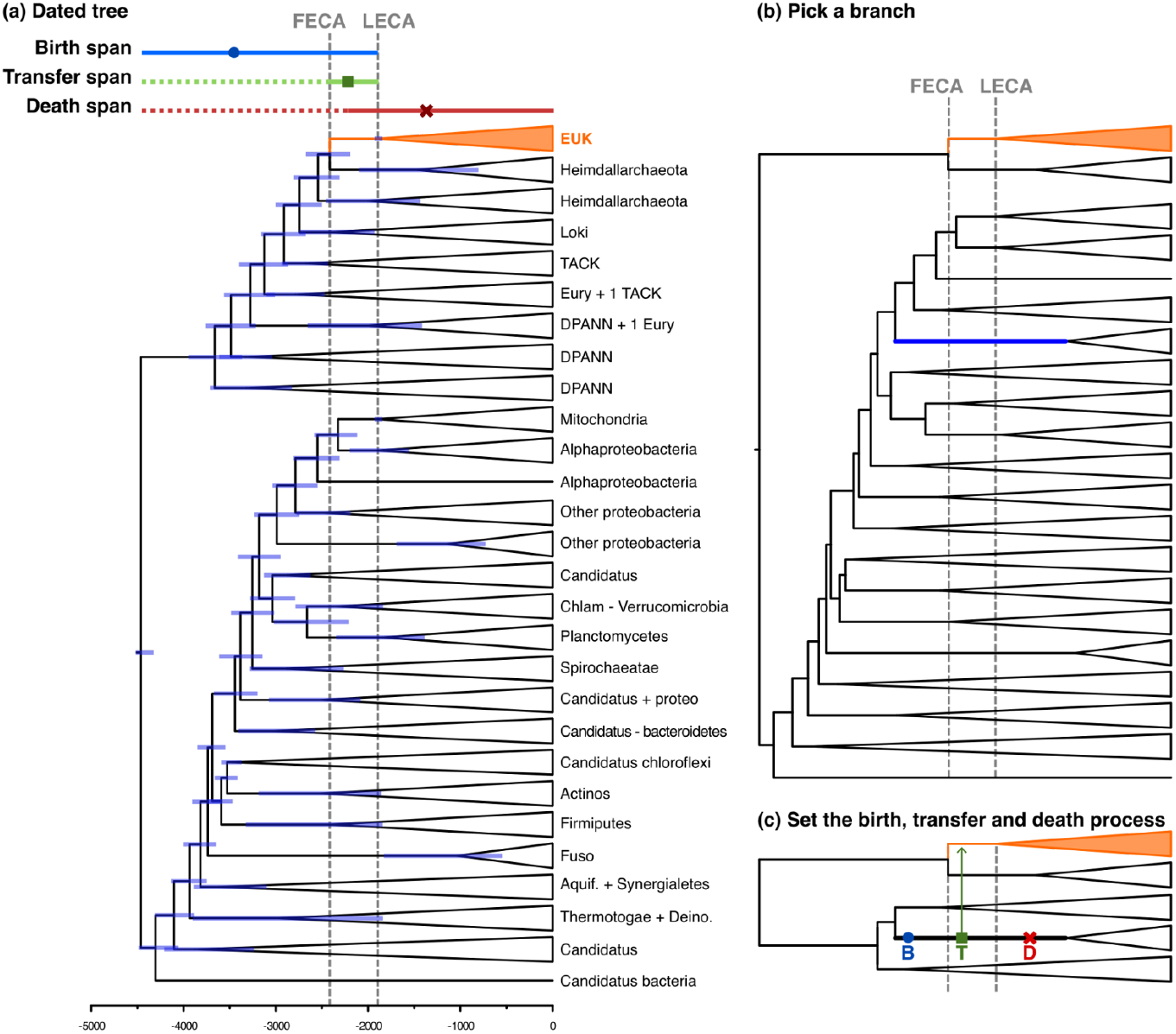
Simulation framework for inferring transfers to the FECA-LECA branch using a dated tree of life. a) Cross-braced dated Tree of Life (Mahendrarajah et al. 2023), solid lines above the tree show the spans used for inferring the events, and the vertical grey dashed lines show the FECA-LECA period. b) Pruned tree showing the archaeal eukaryotic sister and all the bacterial lineages. The thicker blue line is the chosen branch for inferring the ghost. c) The events that are inferred across the chosen branch: the ghost lineage birth (B) the transfer event (T) and the ghost death (D), occurring after the transfer (for further details, see methods).

To simulate transfers from ghost lineages to the proto-eukaryote under the aforementioned premises, we used the following process: (i) we randomly selected a branch from the FECA-LECA branch space (Supplementary Fig. S1), which is then considered the extant sister of the ghosts. (ii) The birth and transfer events of the ghost lineage are inferred depending on the birth and death of the selected (sister) branch: If the branch exists during the whole FECA-LECA period, the ghost birth is calculated between the birth and LECA, and the ghost transfer to the FECA-LECA branch is inferred in the period between the ghost birth and LECA; if the branch originated before FECA and died in the FECA-LECA period, the ghost birth is inferred from the birth and the death of the branch, the transfer is then sampled in the segment of the branch that falls in the FECA-LECA period; if the ghost birth is inferred before FECA, we sampled the transfer from FECA to LECA, whereas when the ghost birth is sampled inside the FECA-LECA period, we inferred the transfer in the period between the ghost birth and LECA; and if the branch originated after FECA and died after LECA, we inferred the ghost birth between the branch birth and LECA, and the transfer between the ghost birth and LECA.

### Conclusion shift test

In a simulation round, we generated 1,000 pairs of transfers from simulated ghosts. For each pair, each one of the transfers was classified as “early” or “late” with respect to the LECA node. We then retrieved the date of the birth of the extant sister clade from within the simulated ghost branches, that is, the date which we would infer in the gene tree. The birth dates of the sisters were then also classified as “early” and “late” with respect to the LECA node, and we, finally, assessed whether the births of the sister clades and the actual transfers had the same order or, otherwise, resulted in a shift, in the latter case, we called a conclusion shift. This simulation round was then repeated 1,000 times (a total of 1,000,000 transfer pairs), allowing us to get a distribution of the percentage of conclusion shifts and the number of branch lengths that, once shifted, result in distances longer than FECA (and that therefore would be considered aberrant and discarded from analysis). We implemented this simulation procedure in R, the code for these functions is stored in the GitHub repository for the project https://github.com/Gabaldonlab/ghost_transfers.

## Results

To assess the likelihood of different putative ghost lineages to mislead the inference of relative timing of gene transfer events during eukaryogenesis using the branch length ratio method (Pittis & Gabaldón 2016a), we used a simulation framework grounded in current knowledge of the Tree of Life and focused on transfers to the relevant lineage: the one spanning from the First Eukaryotic Common Ancestor (FECA) to the Last Eukaryotic Common Ancestor (LECA) (Gabaldón 2021) the FECA-LECA period hereafter. In this framework (see Methods), we simulated ghost lineages and their transfers to the FECA-LECA lineage on top of a recently dated Tree of Life (Fig. 1c). After simulating a million pairs of ghosts and their respective transfers, we calculated the fraction of shifts (i.e. when the inferred order of the transfers is reversed with respect to the simulated -true-one), as well as the fraction of shifts for which one of the transfers is inferred to be older than FECA, and therefore would have not been considered in the original studies. Our results indicate a rather low overall fraction of shifts around 18% (Fig. 2a). That is, around 18% of the gene tree pairs would point to erroneous conclusions in the relative order of transfer events. Conversely, 82% of pairs of gene trees would lead to the right conclusion regarding their relative order, despite the actual donors being ghosts. Moreover, of the 18% of shifts, roughly 18% would result in distances outside FECA-LECA (Fig. 2b), and therefore would not be considered in the context of eukaryogenesis when we can delimit the event. This leads to about 15% of truly confounding gene trees. We believe this to be an acceptable percentage, given other sources of phylogenetic uncertainty (e.g., ortholog inference, evolutionary model assumptions, etc.) (Steenwyk et al. 2023). These trends are similar to those observed when using an alternative dated ToL-scale phylogeny with a more sparse taxonomic sampling (Moody et al. 2024) (the LUCA tree hereafter, Supplementary Fig. S2). In this alternative scenario the proportion of shifted conclusions is similar (21% in average, Supplementary Fig. S2a) than in our main analysis. However, we observe a higher proportion of older than FECA shifts (45%, Supplementary Fig. S2b). This might be caused by the lower taxonomic sampling of the LUCA tree, reduced branch space (Supplementary Fig. S3a and b), as it leads to older births of the clades and ghosts (15% of older births for the ATPase tree over 33% in the LUCA tree, Supplementary Fig. S3c). This distribution of the data causes older estimates of the process in the absence of the closest relatives and, therefore, a bigger proportion of shifted simulations outside the FECA-LECA period. Although we observe this trend, the distributions of relative distance between the ghost birth (the detected age for the transfer) and the actual transfer age are quite similar, suggesting that the simulation process is not driving this difference, but the tree topology and its taxonomic sampling (Supplementary Fig. S3d). Following what we observe in the ATPase tree, there are no conflictive clades that are inferred to have been involved in eukaryogenesis (e.g., Heimdalarchaeia is among the clades with lower pairwise shift probability with the rest of clades, Supplementary Fig. 2c).

**Fig 2.**
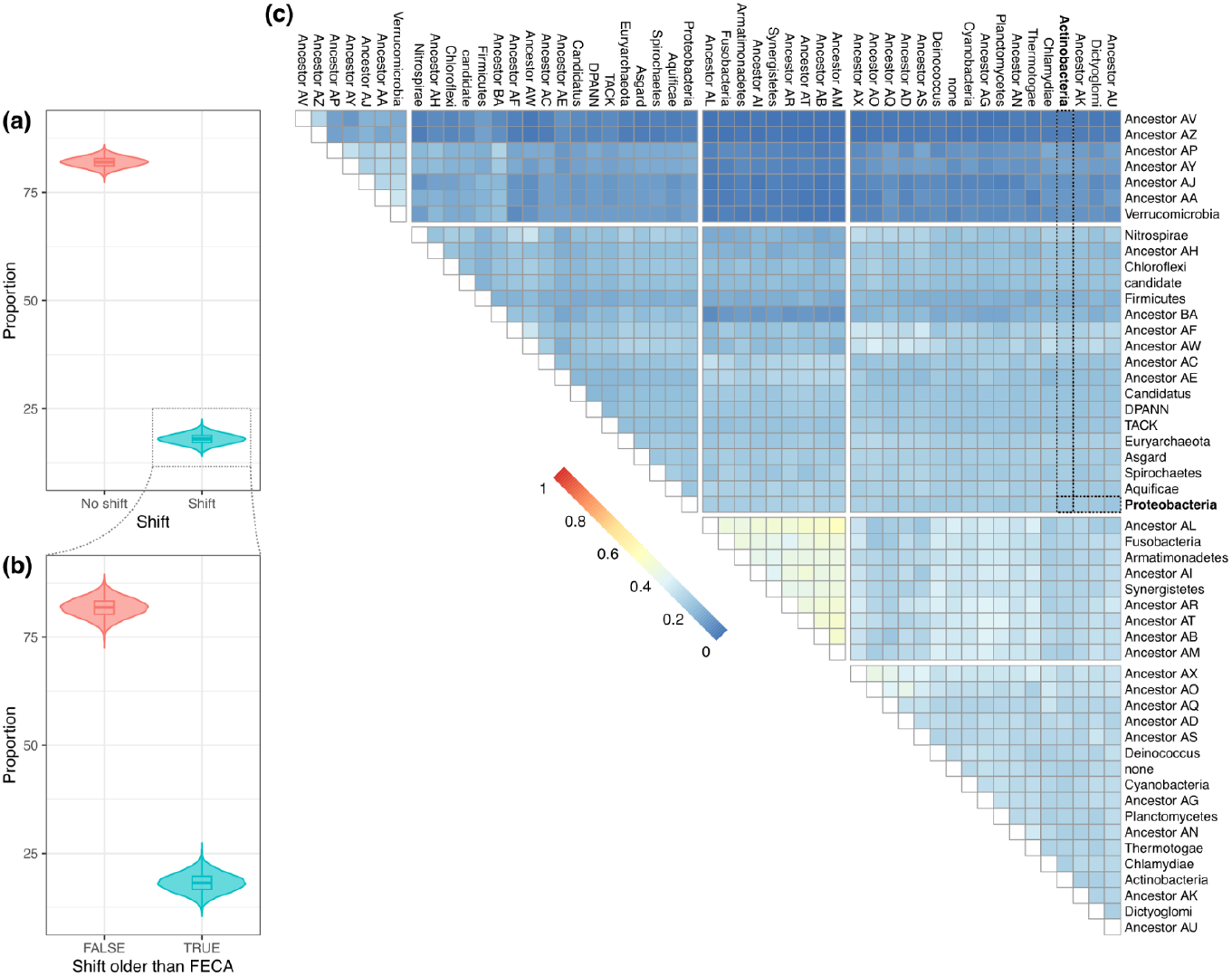
Proportion of shifted conclusions. a) Distributions for the proportion of shifts observed between pairs of simulated ghosts (left). b) For the shift-inducing cases, proportion of those that show an inferred distance older than FECA (outside FECA-LECA period). c) Proportion of shifts per pair of lineages, that is the proportion of simulations resulting in a shift in transfers from the specified pair ancestors. Dashed lines show the relationships of Proteobacteria and Actinobacteria, in bold. The ancestors and their descendant phyla are associated in Supplementary Table S1.

Another advantage of our phylogeny-aware approach is that it allows us to stratify the likelihood of shifts according to the phylogenetic position of the ghost donor, thereby pinpointing clades with particularly high risks of causing shifts. In Pittis and Gabaldón (2016a), each LECA gene family was identified as descending from a prokaryotic origin by attending to the Most Recent Common Ancestor (MRCA) of the sequences in the first prokaryotic sister group. This MRCA could be of any taxonomic level, and therefore, if the prokaryotic sister clade contained sequences from 2 or more phyla (because the transfer occurred prior to the diversification of those phyla, for example), the gene family was assigned to be of uncertain prokaryotic origin and discarded from subsequent analysis. We have decided, however, to analyse the shift likelihood of these lineages, and have employed placeholder names for the ancestors of these bacterial phyla (Supplementary Table S1).

Our results (Fig. 2c) show that some pairs of phyla are more likely to cause shifts than others. Particularly, simulations where ghosts originated from lineages Dictyoglomi (phylum Dictyoglomota *sensu* GTDB), “Ancestor O” (the branch subtending the MRCA of Coprothermobacterota and Caldiserica, see Supplementary Table S1), Planctomycetes (Planctomycetota), Deinococcus (Deinococcota) and Thermotogae (Thermotogota) resulted in a higher proportion of shifts when compared to other phyla, and are particularly problematic when paired with each other. These groups, entailing a higher risk to produce misleading inference when transfers originated from ghost lineages should be taken into account for further analyses of gene flow during eukaryogenesis. Nevertheless, these particular clades are not discussed as potential significant gene donors in previous phylogenomic analysis (Pittis & Gabaldón 2016a; Vosseberg et al. 2021), which reassure their conclusions. Regarding the pair Actinobacteria-Proteobacteria, which was previously analysed in (Pittis & Gabaldón 2016a; Vosseberg et al. 2021), it possesses a moderate-low risk of 27.78% (Fig. 2c), in line with the general results and still indicating that roughly 70% of trees would yield the correct result. Within the 27.78% of shifts, some would yield branches older than FECA (12.7% of those pairs) when analysing a dated tree. Therefore, the actual fraction of confounding shifts would be around 24%.

## Discussion

Incongruence between gene trees and species trees can reveal past events of non-vertical evolution, and the length of the relevant branch underlying the transfer can inform on the relative timing of the event. This information has been the basis of the branch length ratio method for estimating the relative timing of events during eukaryogenesis (Pittis & Gabaldón 2016a; Vosseberg et al. 2021). However, correct inference of such events can hinge on technical factors such as incomplete or unbalanced taxon sampling (Steenwyk et al. 2023; Susko et al. 2021).

The inference of a timeline for the gene transfers to an ancestor is sensitive to the presence of the respective donors and their correct identification in sequence databases. Tricou et al. (2022) assessed the potential effect of incomplete taxon sampling by means of generic tree simulations that were not informed on current knowledge of eukaryogenesis. Here, we added a taxonomic and phylogenetic background to simulate the transfers from prokaryotes to the proto-eukaryote in light of the current knowledge of the Tree of Life. Using this framework, we assessed the proportions of donor clades that, when missing, would lead to erroneous inferences, as well as the impact of alternative species tree topologies when analysing shift probabilities.

Despite a non-negligible contribution of ghost-induced shifts (roughly 20% in average), our results indicate that the vast majority of the trees would point to the right ordering, and that ghosts are unlikely to significantly affect the inference of transfers to the FECA-LECA period. This trend is maintained in a tree with a less dense taxonomic coverage, although with a higher value for ghost-induced shifts. This suggests that a pervasive and abundant taxonomic representation is crucial to reduce the effect of ghost lineages. Moreover, we analysed the pairs of taxa inducing more conclusion shifts. Within shift-inducing transfer pairs, around 20% would cause branches longer than FECA and would then be discarded from subsequent analysis, thereby further reducing the impact of shift-inducing ghosts. However, the detection of branches longer than FECA can only be assessed by using dated trees. Despite this, branches longer than FECA would look like extreme outliers and be discarded on undated trees: an interesting approach would be to assess *modules*, that is, sets of genes inferred to have been acquired at the same relative time; in this sense, everything falling outside this transfer interval could be discarded.

A valuable information obtained from our simulation framework is the varying likelihoods of shifts, depending on the phylogenetic position of ghost donors. In this respect, the top confounding clades are beyond the scope of currently considered bacterial donors in eukaryogenesis (Pittis & Gabaldón 2016a), and the particular clades of interest (namely Proteobacteria and Actinobacteria) present a risk of erroneous shift similar to the average, and a large majority of gene trees would render correct inference of relative order. Although several confounding effects occur when assessing deep phylogenetic hypotheses, the branch length ratio approach focuses on the distribution of the branch lengths. Thus, the hypothesis contrast provides insights into the problem of incomplete sampling as distributions already contain information about the uncertainty of the transfer events and their donors. Ghost lineages are part of this uncertainty. Our results do not only suggest that the overall impact of ghost lineages in the particular problem of gene flow during eukaryogenesis is likely to be low, but they also point to possible control measures. First of all, simulations serve to anticipate donor clades prone to provide potential misleading results, which can be taken into account when interpreting the results. Secondly, our results suggest that discarding gene trees with unusually long branches and selecting groups of genes with similar branch length distributions may alleviate the potentially misleading effects of ghosts.

Altogether our proposed simulation approach clarifies the impact of ghosts in the relative timing of gene transfer during eukaryogenesis and lends additional support to previously made inferences. Moreover, it suggests approaches to identify potential artifactual results and alleviate the confounding effects of ghost lineages. Beyond the relevant study of the origin of eukaryotes, our approach can be easily extended to the study of other evolutionary questions.

## Acknowledgements

TG group acknowledges support from the Spanish Ministry of Science and Innovation (grant numbers PID2021-126067NB-I00, CPP2021-008552, PCI2022-135066-2, and PDC2022-133266-I00), cofounded by ERDF “A way of making Europe”, as well as support from the Catalan Research Agency (AGAUR) (grant number SGR01551), European Union’s Horizon 2020 research and innovation programme (grant number ERC-2016-724173); Gordon and Betty Moore Foundation (grant number GBMF9742); “La Caixa” foundation (grant number LCF/PR/HR21/00737), and Instituto de Salud Carlos III (IMPACT grant IMP/00019 and CIBERINFEC CB21/13/00061-ISCIII-SGEFI/ERDF).

