## Supplementary file for "Phylogeny-aware simulations suggest a low impact of unsampled lineages in the inference of gene flow during eukaryogenesis"

### Supplementary materials

#### *Supplementary figures*

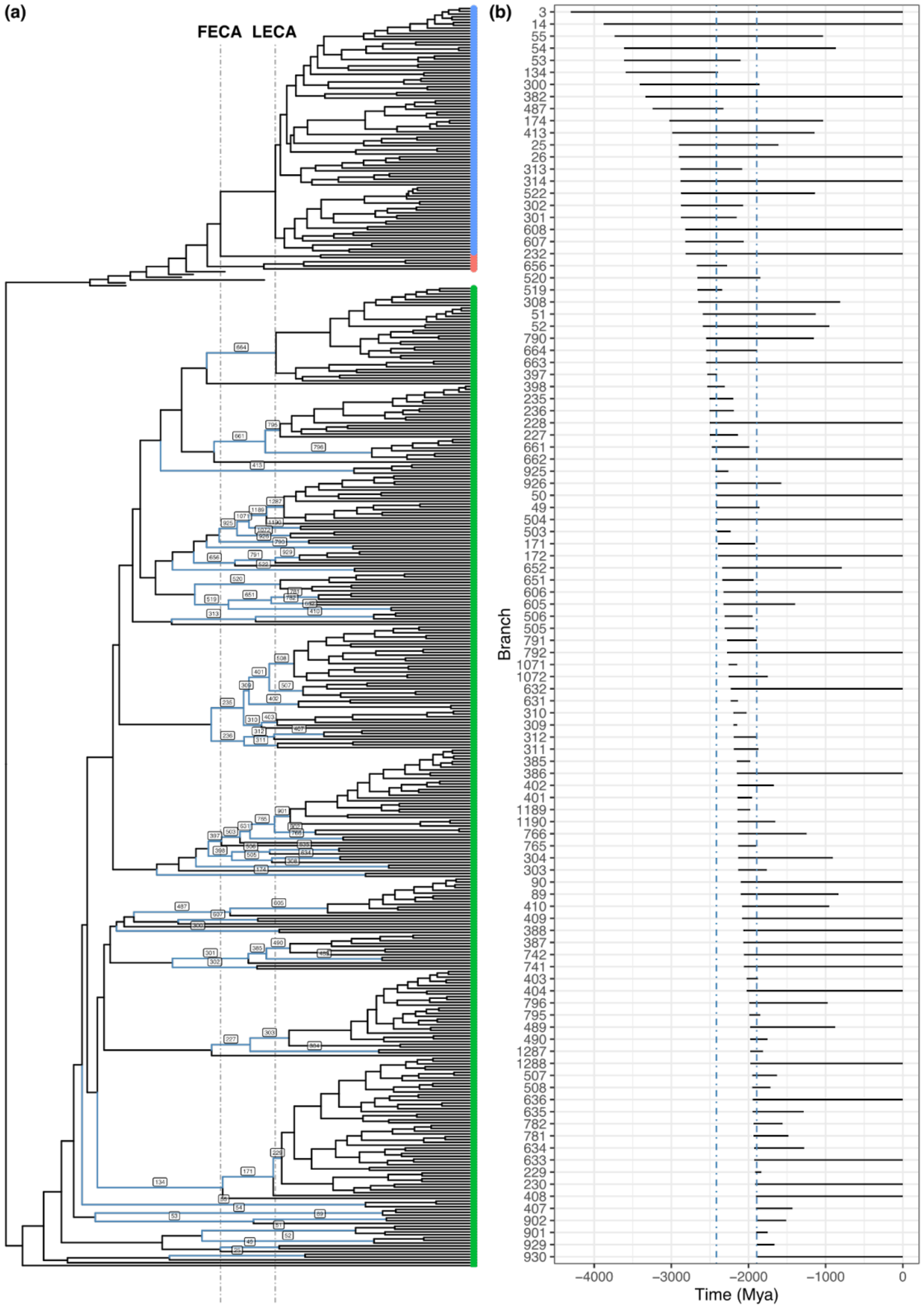

**Supplementary Fig. S1. The branch space**. a) Dated phylogeny from Mahendrarajah et al. (2023) with the branches present in the FECA-LECA period in blue, the branch label corresponds to the number of the branch. Split branches correspond to archaeal lineages different from Asgardarchaeota. b) The set of branches present in the FECA-LECA period sorted by birth date, the number corresponds to the number of the branch in a).

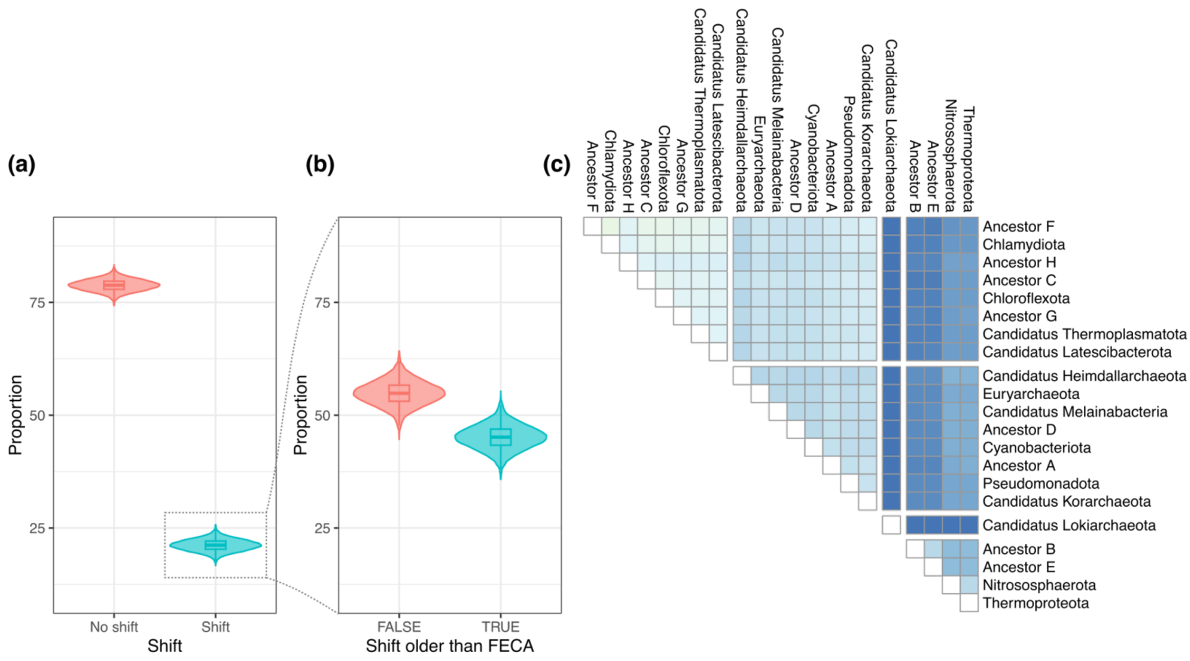

**Supplementary Fig. S2. Proportion of shifted conclusions for the tree from Moody et al.** (2024). a) Distributions for the proportion of shifts observed between pairs of simulated ghosts (left). b) For the shift-inducing cases, proportion of those that show an inferred distance older than FECA (outside FECA-LECA period). b) Proportion of shifts per pair of lineages, that is the proportion of simulations resulting in a shift in transfers from the specified pair ancestors. Pseudomonadota contains the proteobacterial clades. The ancestors and their descendant phyla are associated in Supplementary Table S2.

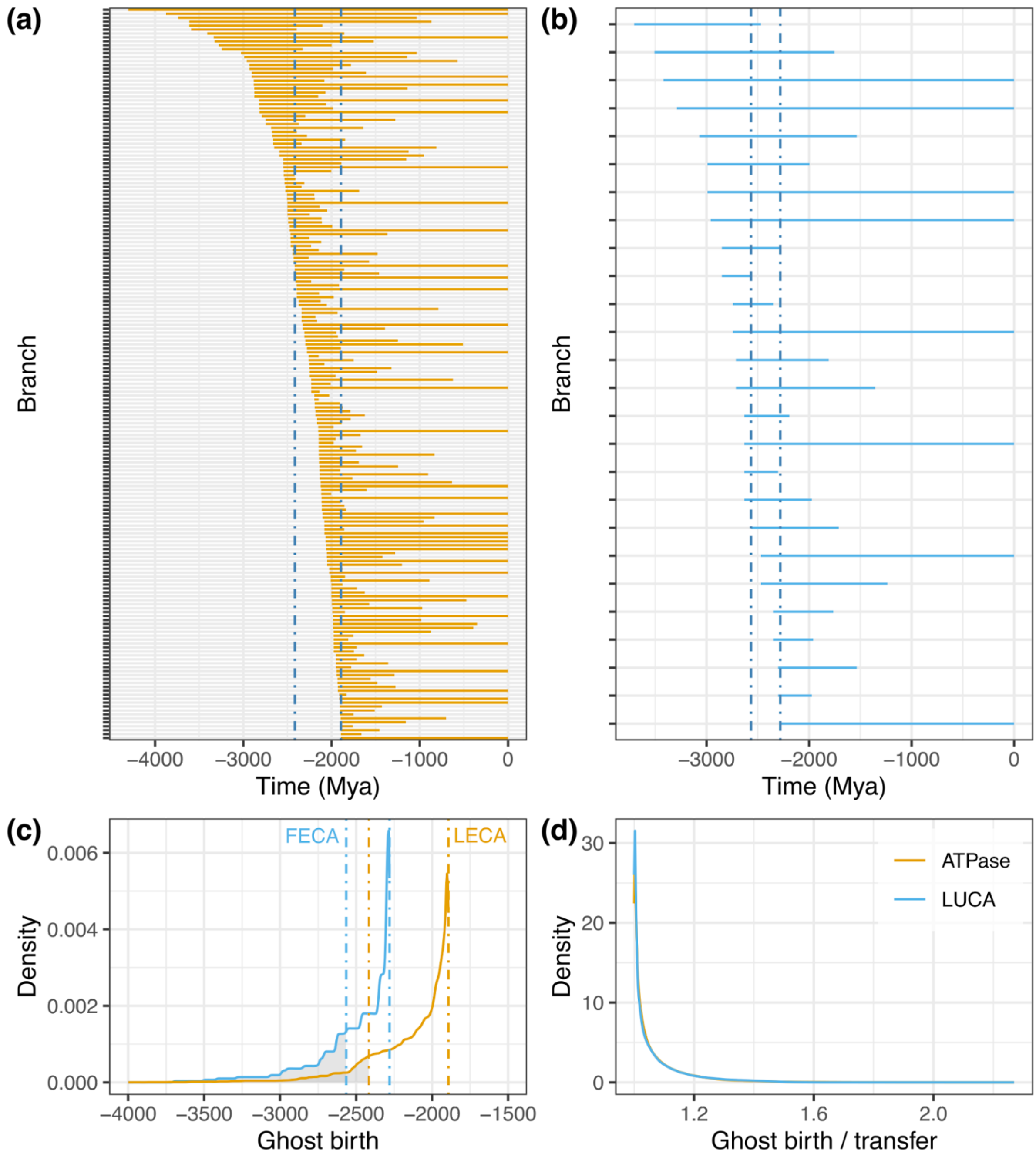

**Supplementary Fig. S3. The effect of sampling in the branch space**. a) ATPase tree (Mahendrarajah et al. 2023) branch space coexisting with LECA. b) LUCA tree (Moody et al. 2024) branch space coexisting with LECA. c) Distribution of the birth of the simulated ghosts, the vertical dashed lines show the FECA and LECA estimates for each tree, and the shadowed area is the proportion of ghost births older than FECA. d) Ratio between the ghost birth and the transfer ages. The colours of the line in all the panels show the trees used following the legend in panel d).

### *Supplementary Tables*

**Supplementary Table S1.** Correspondence between ancestral donor nodes and their extant daughter phyla. The none clade refers to a set of clades that do not have an associated phylum, as the taxonomy is inherited from the original tree in Mahendrarajah et al. (2023).

| Ancestor | Members |
|---|---|
| **Ancestor AA** | Lentisphaerae |
| | Kiritimatiellaeota |
| **Ancestor AB** | Rhodothermaeota |
| | Balneolaeota |
| | Bacteroidetes |
| | Ignavibacteriae |
| | none |
| | Chlorobi |
| | Candidatus |
| **Ancestor AC** | DPANN |
| | Euryarchaeota |
| **Ancestor AD** | Kiritimatiellaeota |
| | Lentisphaerae |
| | Verrucomicrobia |
| **Ancestor AE** | Tenericutes |
| | Firmicutes |
| **Ancestor AF** | Euryarchaeota |
| | TACK |
| **Ancestor AG** | TACK |
| | Archaea |
| **Ancestor AH** | Candidatus |
| | Gemmatimonadetes |
| **Ancestor AI** | candidate |
| | Candidatus |
| **Ancestor AJ** | Fibrobacteres |
| | Candidatus |
| **Ancestor AK** | none |
| | Candidatus |
| **Ancestor AL** | Kiritimatiellaeota |

| | |
|---|---|
| | Chlamydiae |
| | Lentisphaerae |
| | Verrucomicrobia |
| **Ancestor AM** | Fibrobacteres |
| | candidate |
| | Gemmatimonadetes |
| | Candidatus |
| **Ancestor AN** | Elusimicrobia |
| | Candidatus |
| **Ancestor AO** | Proteobacteria |
| | Acidobacteria |
| | Candidatus |
| **Ancestor AP** | Chrysiogenetes |
| | Nitrospinae |
| | Deferribacteres |
| | Candidatus |
| **Ancestor AQ** | Rhodothermaeota |
| | Balneolaeota |
| | Bacteroidetes |
| | Ignavibacteriae |
| | Chlorobi |
| | Candidatus |
| **Ancestor AR** | Proteobacteria |
| | Nitrospinae |
| | Nitrospirae |
| | Chrysiogenetes |
| | Acidobacteria |
| | Deferribacteres |
| | Thermodesulfobacteria |
| | Candidatus |
| **Ancestor AS** | Fibrobacteres |
| | candidate |
| | Candidatus |

| | |
|---|---|
| **Ancestor AT** | TACK |
| | Euryarchaeota |
| **Ancestor AU** | Coprothermobacterota |
| | Caldiserica |
| **Ancestor AV** | Ignavibacteriae |
| | Candidatus |
| **Ancestor AW** | Proteobacteria |
| | Nitrospinae |
| | Chrysiogenetes |
| | Deferribacteres |
| | Thermodesulfobacteria |
| | Candidatus |
| **Ancestor AX** | Proteobacteria |
| | Nitrospinae |
| | Nitrospirae |
| | Chrysiogenetes |
| | Deferribacteres |
| | Thermodesulfobacteria |
| | Candidatus |
| **Ancestor AY** | Proteobacteria |
| | Thermodesulfobacteria |
| **Ancestor AZ** | Rhodothermaeota |
| | Chlorobi |
| | Balneolaeota |
| | Bacteroidetes |
| **Ancestor BA** | Acidobacteria |
| | Candidatus |

47

**Supplementary Table S2.** Correspondence between ancestral donor nodes and their extant daughter phyla. The taxonomy is inherited from the original tree in Moody et al. (2024).

| Ancestor | Members |
|---|---|
| **Ancestor A** | Chlorobiota |
| | Candidatus Cloacimonadota |
| | Elusimicrobiota |
| | Calditrichota |
| | Verrucomicrobiota |
| | Synergistota |
| | Lentisphaerota |
| | Dictyoglomota |
| | Caldisericota |
| | Planctomycetota |
| | Campylobacterota |
| | Thermotogota |
| | Fibrobacterota |
| **Ancestor B** | Chlorobiota |
| | Elusimicrobiota |
| | Calditrichota |
| | Verrucomicrobiota |
| | Synergistota |
| | Lentisphaerota |
| | Planctomycetota |
| | Campylobacterota |
| | Candidatus Cloacimonadota |
| | Fibrobacterota |
| **Ancestor C** | Nitrososphaerota |
| | Thermoproteota |
| **Ancestor D** | Acidobacteriota |
| | Nitrospirota |
| | Thermodesulfobacteriota |
| | Aquificota |
| **Ancestor E** | Thermotogota |
| | Dictyoglomota |

| | |
|---|---|
| | Caldisericota |
| **Ancestor F** | Bacillota |
| | Actinomycetota |
| **Ancestor G** | Euryarchaeota |
| | Candidatus Thermoplasmatota |
| **Ancestor H** | Candidatus Thorarchaeota |
| | Candidatus Lokiarchaeota |
| | Candidatus Odinarchaeota |

***Supplementary references***

Mahendrarajah TA et al. 2023. ATP synthase evolution on a cross-braced dated tree of life. Nat Commun. 14:7456. doi: 10.1038/s41467-023-42924-w.

Moody ERR et al. 2024. The nature of the last universal common ancestor and its impact on the early Earth system. Nat Ecol Evol. 8:1654–1666. doi: 10.1038/s41559-024-02461-1.
